# Towards Inferring Nanopore Sequencing Ionic Currents from Nucleotide Chemical Structures

**DOI:** 10.1101/2020.11.30.404947

**Authors:** Hongxu Ding, Ioannis Anastopoulos, Andrew D. Bailey, Joshua Stuart, Benedict Paten

**Affiliations:** Department of Biomolecular Engineering, UC Santa Cruz, Santa Cruz, California, USA; UC Santa Cruz Genomics Institute, Santa Cruz, California, USA

## Abstract

The characteristic ionic currents of nucleotide kmers are commonly used in analyzing nanopore sequencing readouts. We present a graph convolutional network-based deep learning framework for predicting kmer characteristic ionic currents from corresponding chemical structures. We show such a framework can generalize the chemical information of the 5-methyl group from thymine to cytosine by correctly predicting 5-methylcytosine-containing DNA 6mers, thus shedding light on the *de novo* detection of nucleotide modifications.

## INTRODUCTION

During nanopore sequencing, consecutive nucleotide sequence kmers block the pores sequentially, producing ionic currents [1]. Chemical modifications on nucleotides additionally alter the ionic currents measured during nanopore sequencing [2–9]. The characteristic ionic currents of kmers, which are represented in kmer models, are used in interpreting nucleotide modifications [2–5]. Up to now, 7 [2–4, 6–8] and 4 [5,9] modifications have been successfully characterized in DNA and RNA, respectively. To date, most modification prediction algorithms are based on kmer models [2–4,6]. However, such learning strategies struggle to generalize knowledge between related kmers. For example, our previous hierarchical Dirichlet process approach could be structured to learn associations between kmers with specific shared properties, e.g. by numbers of pyrimidine bases, but could not generally learn relationships between arbitrary chemical similarities [2]. Moreover, such approaches necessarily represent base modifications as distinct, unrelated characters. The upshot being that such kmer character-based models require extensive training data and are unable to *de novo* predict the impact of a chemical modification. Given that the number of possible kmers increases polynomially with the number of modifications being modeled, it is extremely challenging to generate sufficient control data for such models, especially considering that more than 50 and 160 nucleotide modifications have been verified in DNA and RNA respectively [10,11].

To start to tackle this problem, we propose a graph convolutional network (GCN)-based deep learning framework [12,13] for predicting kmer characteristic ionic currents from corresponding kmer chemical structures. We confirm that the proposed framework is able to represent individual kmer chemical modules, such as the phosphate group, the sugar backbone, as well as the nucleobase methyl and amine groups. We further demonstrate that this framework can infer full kmer models even when the training data does not include all possible kmers. This opens up the possibility of modeling kmers that are under-represented in control datasets. We also show the framework can generalize the 5-methyl group in thymine to cytosine, thereby accurately predicting the characteristic ionic currents of 5-methylcytosine (5mC)-containing DNA 6mers. Such generalization of chemical information is a reason for optimism about the potential for *de novo* detection of nucleotide modifications.

## RESULTS

### Architecture of the deep learning framework

Our deep learning framework consists of three groups of layers, including GCN layers, convolutional neural network (CNN) layers, and one fully connected neural network (NN) layer. As shown in Figure 1A, the kmer chemical structures are first represented as graphs, with atoms as nodes and covalent bonds as edges. The atom chemical properties are then assigned as node attributes. Based on such graphs, GCN layers extract one chemical feature vector for every atom, by visiting its immediate graph neighbors. By this means, after several GCN layers, atom feature vectors will contain chemical information for all atoms within a certain graph distance. Specifically, this distance equals the number of GCN layers applied. Considering the small encoding distance of each layer of a GCN, to improve the encoding efficiency of the framework, CNN layers are then applied as an optimization to summarize relatively long-range chemical information above the GCN layers. The output matrices of the final CNN layer are then “flattened” as feature vectors. Such feature vectors are then passed to the final fully connected NN layer to summarize kmer-level information and finally predict the kmer characteristic ionic currents (see METHODS). For DNA and RNA, the corresponding best-performing architecture in hyper-parameter tuning was selected for downstream analysis (see METHODS).

**Figure 1.**
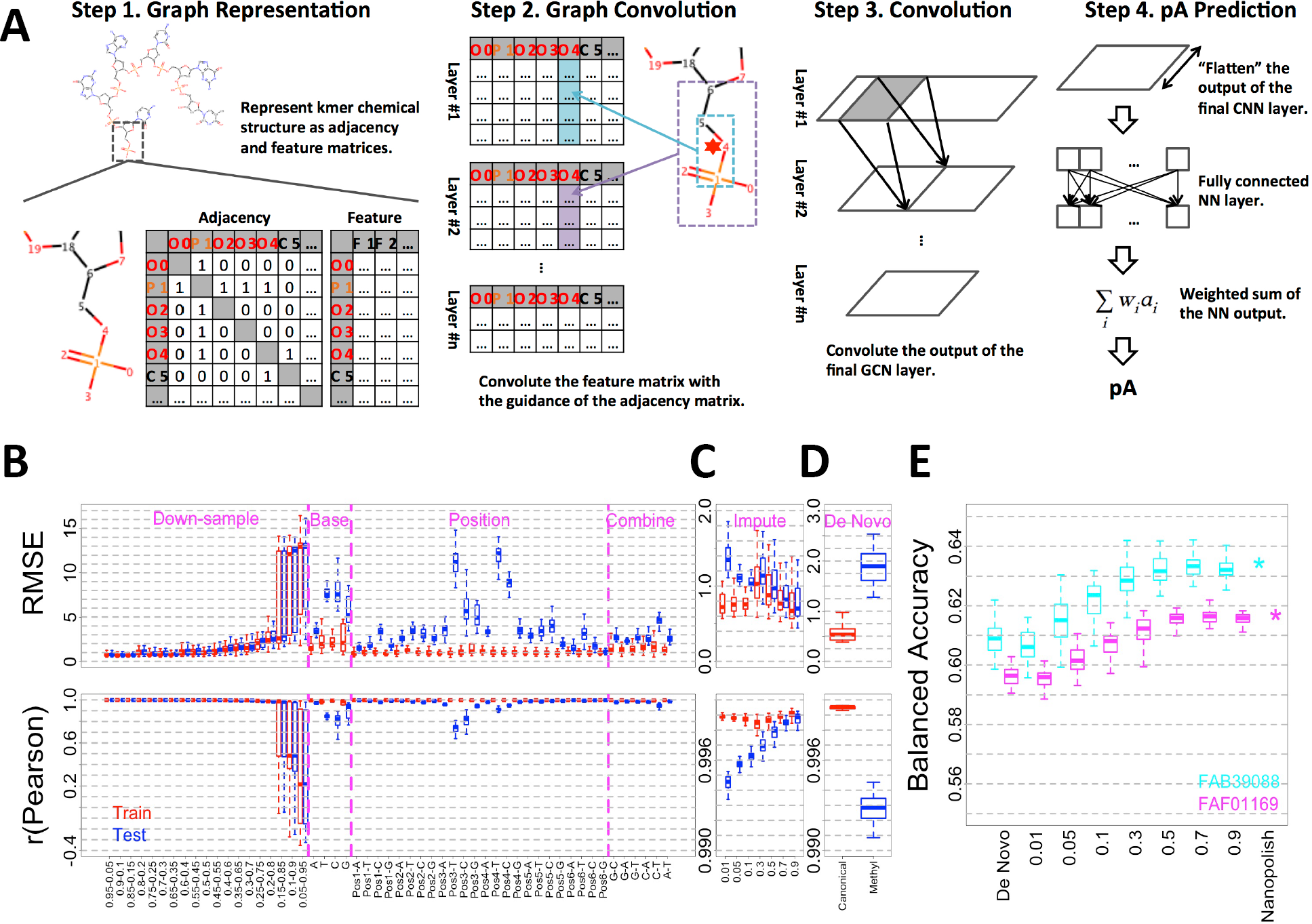
Predicting kmer characteristic ionic currents from chemical structures. (A) Graphic overview of the proposed deep learning framework for DNA analysis. (B) Goodness-of-fit of DNA canonical random down-sample, base-dropout, position-dropout and model combination analyses. (C) Goodness-of-fit of 5mC-containing DNA 6mer imputation analysis. (D) Goodness-of-fit of *de novo* 5mC-containing DNA 6mer prediction. In panel B-D, Train (red) and Test (blue) refer to goodness-of-fit of the training and test DNA 6mers, respectively. (E) Predictive accuracy of C-5mC status quantified by balanced accuracy. Nanopolish, nanopolish model as baseline control. De Novo, 5mC-containing DNA 6mer models predicted from canonical training. 0.01-0.9, different imputation fractions of 5mC-containing DNA 6mers. FAB39088 (cyan) and FAF01164 (purple) refer to two independent NA12878 cell line native genomic DNA nanopore sequencing datasets.

### Kmer-level generalization

We first confirmed the proposed framework can accurately predict characteristic ionic currents of kmers from their chemical structures. To do so, we performed down-sample analysis on the canonical DNA 6mer model provided by Oxford Nanopore Technologies (ONT, see METHODS), by randomly partitioning canonical DNA 6mers with various train-test splits. For each train-test split group, we performed 50-fold cross validation, and used root mean square error (RMSE) and Pearson correlation (r) to quantify the goodness-of-fit (see METHODS). As shown in Figure 1B and Supplementary Figure 1, the performance stabilized as more than 40% of DNA 6mers were included in the training. Specifically, for DNA 6mers only used in the test, average RMSE and Pearson correlation reached 1 and 0.995, respectively. Such a result indicated on average 40% of randomly selected DNA 6mers contain sufficient information to recapitulate the full DNA 6mer model.

We next explored how training specific kmer subsets influence the ionic current predictions. Specifically, we trained the framework using either the DNA 6mers that **a)** do not contain a given nucleotide (base-dropout), **b)** do not specify a nucleotide at a given position (position-dropout) and **c)** that are combined from different base-dropouts (for instance using the union of A-dropout and T-dropout kmers, such that kmers containing both A and T would be excluded, but not kmers containing either A or T, noted as A-T model combination, see METHODS and Supplementary Note 1 for details). This latter combination analysis simulates the situation in which we have knowledge about two modifications independently, but must guess at the effect of their combination. For each group in **a-c)** 50 independent repeats were performed, and goodness-of-fit was used to evaluate the performance. As shown in Figure 1B and Supplementary Figure 1, base and position-dropouts significantly decreased the prediction power. Moreover, dropouts in 3rd and 4th positions contributed the most to prediction power decrease, followed by 2nd and 5th positions, consistent with [14]. Model combinations, on the other hand, in general had a minor influence on the prediction power.

The above-mentioned analyses together suggest, once properly trained with sufficient and diverse 6mers, the kmer-level generalizability of the framework. To further validate and extend our framework we performed all the above-mentioned analyses under RNA context. Considering the significantly smaller amount of training data (1/4th the number of distinct RNA 5mers vs DNA 6mers), the prediction power of the RNA architecture is compromised. However, once trained with a similar number of kmers, the RNA architecture yielded comparable prediction power. For instance, the RNA 0.95-0.05 (972 training kmers) and DNA 0.25-0.75 (1024 training kmers) train-test splits yielded comparable performance on test data. Such a result suggests the validity of our proposed architecture (see METHODS, Supplementary Figure 2 and Supplementary Note 2 for details).

**Figure 2.**
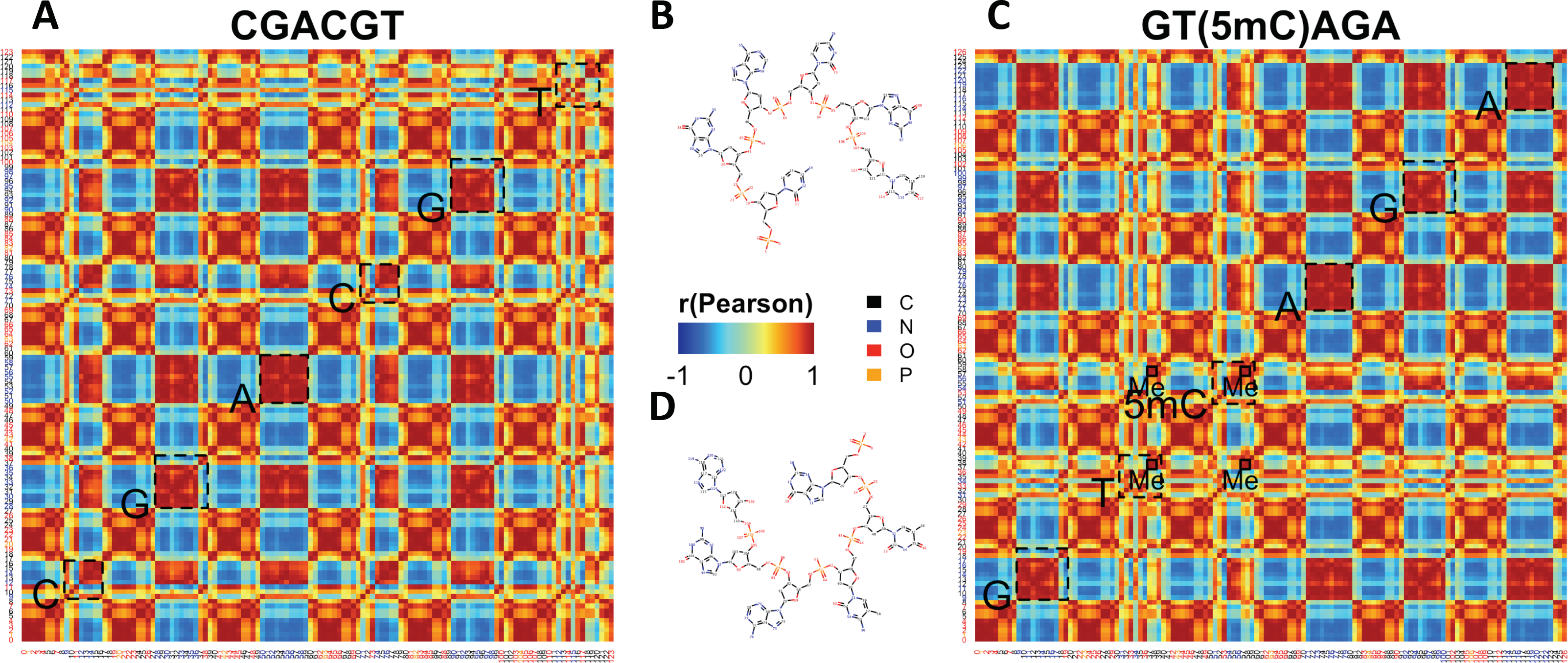
Visualizing the encoding of chemical structures. (A, B) Chemical structure and atom similarity matrix of example canonical DNA 6mer CGACGT. Nucleobases were highlighted by dashed boxes. Atoms were numbered and colored based on the chemical structure in (B). Carbon, nitrogen, oxygen and phosphorus were colored as black, blue, red and orange, respectively. (C, D) Chemical structure and atom similarity matrix of example 5mC-containing DNA 6mer GT(5mC)AGA. Specifically, methyl groups (Me) in T and 5mC were highlighted in solid boxes. Atoms were numbered and colored based on the chemical structure in (D). Carbon, nitrogen, oxygen and phosphorus were colored as black, blue, red and orange, respectively.

Such kmer-level generalizability could facilitate nucleotide modification detection by greatly reducing the required control data to generate reliable full modification-containing kmer models. As a proof-of-concept, we trained the DNA deep learning architecture with all canonical 6mers plus {1%, 5%, 10%, 30%, 50%, 70%, 90%} of randomly selected 5mC-containing 6mers (“modification imputation” analysis). The characteristic ionic current signals of such 5mC-containing DNA 6mers were obtained from the nanopolish model as reported in [3,6]. For each training group 50 independent repeats were performed (see METHODS). As shown in Figure 1C and Supplementary Figure 3, decent goodness-of-fit could be obtained when as few as 5% of 5mC-containing DNA 6mers were used as training data. Specifically, for test DNA 6mers, average RMSE and Pearson correlation reached 1.2 and 0.995, respectively. Furthermore, models trained with knowledge of 50% 5mC-containing DNA 6mers performed about as well as models trained with 90%.

### Chemical group-level generalization and de novo prediction

We noted that performance of the model on held out 5mC kmers trained with just 1% of 5mC kmers was better than chance. This raised the question of if chemical group-level information was being usefully generalized among nucleotides by our framework, potentially allowing the 5mC to be predicted *de novo,* without ever having been seen by the model. As a chemical derivative of cytosine, 5mC contains an additional methyl group at the 5th position (5-methyl) of the pyrimidine ring. This 5-methyl group is shared between 5mC and thymine. We thus hypothesized that 5mC can be generalized by combining the pyrimidine ring from cytosine and 5-methyl group from thymine. As a proof-of-concept, we trained the framework with all canonical DNA 6mers to make *de novo* predictions on 5mC-containing DNA 6mers. Similar to previous analyses, 50 independent repeats were performed, and the prediction power was first quantified by goodness-of-fit against the above-mentioned nanopolish model. As shown in Figure 1D and Supplementary Figure 3, although goodness-of-fit of 5mC-containing DNA 6mers were significantly worse than canonical counterparts, decent performance could still be obtained (average RMSE and Pearson correlation reached 1.8 and 0.993, respectively). We also compared the goodness-of-fit between canonical and 5mC-containing DNA 6mers, and as shown in Supplementary Figure 4, a strong positive correlation could be observed. Such a result confirmed that no overfitting was introduced during architecture-training with canonical DNA 6mers, and further suggested 5-methyl generalization.

### C/5mC status predictive analysis

We next performed “predictive analysis” to test whether the DNA 6mer models inferred by our deep learning framework could be used to correctly predict DNA C/5mC status at a per-read, per-site resolution from ionic currents (“predictive accuracy”, see METHODS). C/5mC-sites to be predicted were confirmed by bisulfite sequencing (see METHODS). We also quantified the predictive accuracy with the above-mentioned nanopolish model as a baseline control (see METHODS). As shown in Figure 1E, average predictive accuracy, quantified by balanced accuracy, became comparable with baseline control with 50% of imputed 5mC-containing 6mers. Taken together, these results confirmed the kmer-level generalizability of our framework, as well as suggesting that reliable modification-containing kmer models can be built with significantly less control data once facilitated by our methodology. Even for the *de novo* case (model trained with 0% of 5mC kmers) balanced accuracy values reached 60%. Such balanced accuracy values were comparable to the baseline control. Such a result confirmed the successful 5-methyl generalization. More confusion matrix-based prediction evaluations can be found in Supplementary Figure 5.

### The encoding of chemical structures

To better understand how chemical structures were encoded we visualized DNA 6mer atom similarity matrices. Specifically, we trained the proposed framework with all canonical DNA 6mers. We then calculated and visualized the Pearson correlations of the feature vectors derived by the final GCN layer as atom-level similarities. As shown in Supplementary Figure 6, we visualized 10 randomly chosen canonical DNA 6mers. Taking CGACGT as an example, as shown in Figure 2A and B, atoms were in general aggregated by chemical contexts. For instance, for the first cytidine monophosphate in CGACGT, atoms #0-4 were tightly clustered with average r>0.9, recapitulating the phosphate group. Atoms #5-8 and #17-18 also clustered with average r>0.9, denoting the deoxyribose backbone. Among cytosine atoms #9-16, #9 nitrogen atom connected the nucleobase to the deoxyribose backbone, atoms #10-11 denoted the C=O group, and atoms #12-16 composed the C=C-C=N conjugation system and the covalently bonded amine group. Similarly, atoms in other nucleotides can also be clustered into phosphate groups, deoxyribose backbones and nucleobases. Within the nucleobases, chemical modules including chemical groups and conjugation systems can further be dissected. Such a phosphate-deoxyribose-nucleobase pattern repeated and constituted DNA 6mers.

We also examined the inter-nucleotide similarities of different components. As shown in Figure 2A and B, in general high similarities (average r>0.9) were observed among phosphates, as well as deoxyriboses from different nucleotides. Meanwhile, chemical modules sharing similar structures, e.g. the conjugation systems of adenines, cytosines and guanines were more similar to each other. On the other hand, low similarities (average r<0.5) were observed between chemical modules with distinct structures, e.g. the cytosine C=O group and the thymine methyl group. Taken together, these results suggest that the GCN layers in the proposed framework can effectively capture features interpretable as individual chemical modules.

We further visualized the atom-level similarity matrices of 5mC-containing DNA 6mers, aiming to understand the generalization of methyl group among thymine and 5-methylcytosine. Following the same above-mentioned process, we generated and visualized the atom-level similarity matrices of 10 randomly selected 5mC-containing DNA 6mers (Supplementary Figure 7). Taking GT(5mC)AGA as an example (Figure 2C and D), the phosphate-deoxyribose-nucleobase repetitive pattern was recapitulated. Within nucleobases, high similarities (average r>0.9) were again observed among chemical modules with similar structures. Specifically, strong similarities (average r>0.9) were observed between thymine (#37-38) and 5mC (#57-58) methyl groups (Me). In addition, such methyl groups were uniquely encoded as they were less correlated with any other DNA 6mer chemical modules (average r<0.5). These observations together suggested the successful chemical information generalization. Noticeably, the methyl groups were encoded with the pyrimidine backbone C=C modules. Such a result suggests that the GCN-encoding is driven by chemical context, which further implies when generalizing one specific chemical group among different nucleotides, the corresponding chemical contexts in which such chemical group resides should be the same.

## DISCUSSION

We propose a GCN-based deep learning framework for associating kmer chemical structures with corresponding characteristic ionic currents. We show that such a framework can recapitulate full kmer models from partial training data, thus greatly facilitating modification analysis by reducing the amount of required control data. Specifically, for cases where a small proportion of random kmers are under-represented in control data, we can apply the same principle as the down-sample analysis to learn around these training deficiencies. For cases where comprehensive control datasets are available only for single modifications, we could apply model combination (as we showed for individual nucleotides) to model kmers containing multiple modifications simultaneously.

We also demonstrated that such a framework can represent novel modifications by generalizing encoded chemical groups between nucleotides, thus shedding light on *de novo* modification detection. Please note that the proposed framework encodes chemical groups, e.g. the methyl groups in thymine and 5mC, as well as the amine groups in cytosine, guanine and adenine, with covalently bonded “backbone atoms”, showing a strong chemical context-specificity (Figure 2 and Supplementary Figure 6, 7). Thus, the current framework cannot properly handle “stacked” chemical groups. For instance, the methylamine group in N6-methyladenine (6mA) cannot be correctly encoded by simply stacking methyl with amine. As shown in Supplementary Figure 8, substituting A with 6mA was predicted to decrease characteristic ionic currents, which is opposite of a previous study [15]. Therefore the extensibility of the framework is largely limited. To overcome such a limitation, controlled nanopore sequencing profiles of diverse nucleotide modifications are needed.

Deep learning-based approaches have emerged as powerful tools for detecting nucleotide modifications from nanopore sequencing readouts. Compared to kmer model-based counterparts, deep learning-based approaches are reported to have better accuracy and less computational resource consumption [7,8]. Thus, one potential future extension of the paper would be using the learned models as components of a larger, recurrent deep neural network.

Another potential future direction would be generalizing the proposed framework to handle both DNA and RNA kmers. Due to different translocation speed, the nanopore sequencing ionic currents of DNA and RNA are not directly comparable [16]. Therefore, advanced deep learning frameworks which can take both kmer chemical structures and nanopore sequencing experimental setups are needed. Considering DNA and RNA share several non-canonical nucleobases, e.g. Inosine (I) [17], we might combine the ribose in RNA and I in DNA to reconstruct I-containing RNA 5mers, and vice versa for I-containing DNA 6mers. By this means, required RNA control nanopore sequencing reads, which are usually challenging to obtain, can be largely compensated. Meanwhile, such generalization would largely diversify the chemical contexts that can be represented, further facilitating the *de novo* modification analysis.

## METHODS

### Methods summary

The deep learning framework proposed here aims to associate kmer chemical structures with corresponding characteristic ionic currents. The chemical structure refers to the chemical properties of kmer atoms and how these atoms are covalently bonded. Characteristic ionic current, on the other hand, refers to the average ionic current that a specific kmer produces during nanopore sequencing.

Thus, in the following sections, we first describe how the chemical structures were represented (“graph representation of kmer chemical structures”). We then describe the deep learning framework used in the study (“architecture of the deep learning framework”, “training procedure” and “hyper-parameter tuning”). We further describe analyses performed to evaluate the performance of the proposed framework (“down-sample, base-dropout, position-dropout and combination analysis”, “predicting modification-containing DNA 6mers” and “predictive analysis of predicted kmer models”). Finally we describe all required resources for the study (“kmer models”, “data availability” and “code availability”).

### Graph representation of kmer chemical structures

Following the workflow described in [12], kmer chemical structures were first described by SMILES (Simplified Molecular Input Line Entry System) strings, which were assembled by concatenating SMILES strings of individual nucleotides, as summarized in the following table. Each nucleotide base can be described by several SMILES strings. The SMILES strings presented in the table below were selected due to the ease of combining them into complete kmers. Based on information provided by Oxford Nanopore Technologies, as well as a previous study [14], DNA and RNA is represented by 6mer and 5mer, respectively. An “O” was then added to the end of each concatenation to represent the residual unbonded hydroxyl group on the sugar backbone.

| Nucleotide | SMILES string |
| --- | --- |
| A (DNA) | OP(=O)(O)OCC1OC(N3C=NC2=C(N)N=CN=C23)CC1 |
| T (DNA) | OP(=O)(O)OCC1OC(N2C(=O)NC(=O)C(C)=C2)CC1 |
| C (DNA) | OP(=O)(O)OCC1OC(N2C(=O)N=C(N)C=C2)CC1 |
| G (DNA) | OP(=O)(O)OCC1OC(N2C=NC3=C2N=C(N)NC3=O)CC1 |
| 5mC (DNA) | OP(=O)(O)OCC1OC(N2C(=O)N=C(N)C(C)=C2)CC1 |
| 6mA (DNA) | OP(=O)(O)OCC1OC(N3C=NC2=C(NC)N=CN=C23)CC1 |
| A (RNA) | OP(=O)(O)OCC1OC(N3C=NC2=C(N)N=CN=C23)C(O)C1 |
| U (RNA) | OP(=O)(O)OCC1OC(N2C(=O)NC(=O)C=C2)C(O)C1 |
| C (RNA) | OP(=O)(O)OCC1OC(N2C(=O)N=C(N)C=C2)C(O)C1 |
| G (RNA) | OP(=O)(O)OCC1OC(N2C=NC3=C2N=C(N)NC3=O)C(O)C1 |

We then represent the SMILES string of each kmer as a graph noted as **G**(**A**, **X**). Specifically, the topology (atom order is determined by SMILES string) of each kmer chemical structure was represented by an adjacency matrix **A**, with **A**_**i,j**_ equals 1 iff the ith and jth atoms were covalently bonded. Meanwhile, for every atom in **A**, the corresponding chemical properties were represented by feature matrix **X**, with **X**_**i**_ recording the chemical property vector for the ith atom. Atom chemical properties included in the study were summarized in the following table:

| Feature | Description |
| --- | --- |
| Carbon | 1 if the atom is carbon, 0 otherwise (boolean) |
| Nitrogen | 1 if the atom is nitrogen, 0 otherwise (boolean) |
| Oxygen | 1 if the atom is oxygen, 0 otherwise (boolean) |
| Phosphorus | 1 if the atom is phosphorus, 0 otherwise (boolean) |
| Atom degree | Total number of covalent bonds around an atom (integer) |
| Implicit valence | It equals the valence of the atom minus the valence calculated from the bond connections (integer) |
| Number of hydrogens | Total count of hydrogens (integer) |
| Aromaticity | 1 if atom in an aromatic ring, 0 otherwise (boolean) |

Therefore, the GCN has encoded as input a chemical feature matrix **X** with the guide of chemical topology matrix **A**, representing kmer chemical structures. Notably, for convenient GCN implementation, the size of **A** and **X** is kept constant. Due to the variable number of atoms across kmers, **A** and **X** were thereby padded with zeros based on the largest kmers. Specifically, the **A** matrix was padded at the end of its rows and columns, with dim(**A**) is {133, 133} and {116, 116} for DNA and RNA, respectively.

While the **X** matrix was padded at the end of its rows, with dim(**X**) is {133, 8} and {116, 8} for DNA and RNA, respectively. Note that the kmer representation is guided by the non-zero elements (covalent bonds) in **A**, thus such padding will not affect the GCN encoding.

### Architecture of the deep learning framework

The Graph Convolutional Network (GCN) layers of our framework were built based on the procedure described by [12]. Fast approximate convolutions on **G** were used to create a graph-based neural network *f*(**X, A**), following the propagation rule:

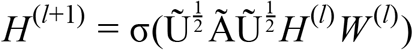

σ(•) is the activation function applied to each layer. Here, the activation function used was Exponential Linear Unit (ELU). 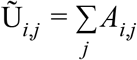 is the degree matrix for each atom in the graph. 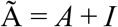 adds self edges to each of the atoms. The 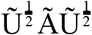 transformation prevents changes in the scale of the feature vectors [13] and constructs filters for the averaging of neighboring node features. **H** and **W** denote the output (activation vectors) and weights of each GCN layer, respectively. The corresponding superscript represents the layer index. **H**^0^ = **X**, however subsequent **H** represent the GCN derived features.

The intuition of the graph convolution process is described as follows. For every kmer, chemical properties of atoms, together with their covalently bonded neighbors, will be convoluted with the guidance of **G**. Such graph convolution yields an activation matrix **H**, following the aforementioned propagation rule. **H** is an atom-by-feature matrix, with dimension {133, N} and {116, N} for each of the DNA and RNA kmers, respectively. Here N equals the number of nodes of the GCN layer, which determines the number of features to be derived. The selection rule for N is described in the following section. As more GCN layers are stacked, the graph convolution process is repeated. The **H** matrix will thus contain chemical information of all atoms within a certain graph distance, which equals the number of GCN layers applied. By this means, “chemical modules” composed of several atoms linked by covalent bonds are encoded.

Considering the small encoding distance of a GCN, for a better encoding efficiency we wanted additional layers that can quickly summarize atom information. We thus applied standard 1-D CNN layers with Rectified Linear Unit (ReLU) activation right after the GCN layers. Average Pooling [18] was applied on the output of each 1-D CNN layer. Average Pooling takes the average of each 2×2 patch of the CNN output matrix. Specifically, output dimension of the first CNN layer equals {133-K+1, N’} and {116-K+1, N’} for DNA and RNA kmers, respectively. Here K is the CNN kernel size and N’ is the node number of the final GCN layer. Output dimensions of subsequent CNN layers equals {m-K+1-2+1, n-2+1}, where {m, n} denotes the output dimension of the previous layer, and 2 denotes the Average Pooling patch size. The output from the final 1-D CNN layer, after Average Pooling, was passed to a Flatten layer, which converts the final 1-D CNN output matrix to a 1-D feature vector in a row-wise fashion. The NN layer then takes the flattened vector as input, thereby summarizing information about the entire kmer, and producing a highly informative representation. Elements of the NN layer output vector are linearly combined as the final pA value.

### Training procedure

Our framework was trained with the Keras [19] framework with TensorFlow backend using the Adam [20] optimizer for gradient descent optimization. The framework was allowed to train for a maximum of 500 epochs. To control for overfitting, EarlyStopping [21] was used by monitoring the increase in validation loss. Early termination of training was reached if the validation loss was increasing for 10 consecutive epochs, indicating that the framework had reached maximum convergence. Mean Squared Error (MSE) was used as the loss function during the training process. Meanwhile, a 10% random dropout was applied after each layer, to further prevent overfitting [22]. In the following experiments the exact same training routine was used.

### Hyper-parameter tuning

In order to determine the optimal architecture, we performed hyper-parameter grid search. The search involved the hyper-parameters shown in the following table:

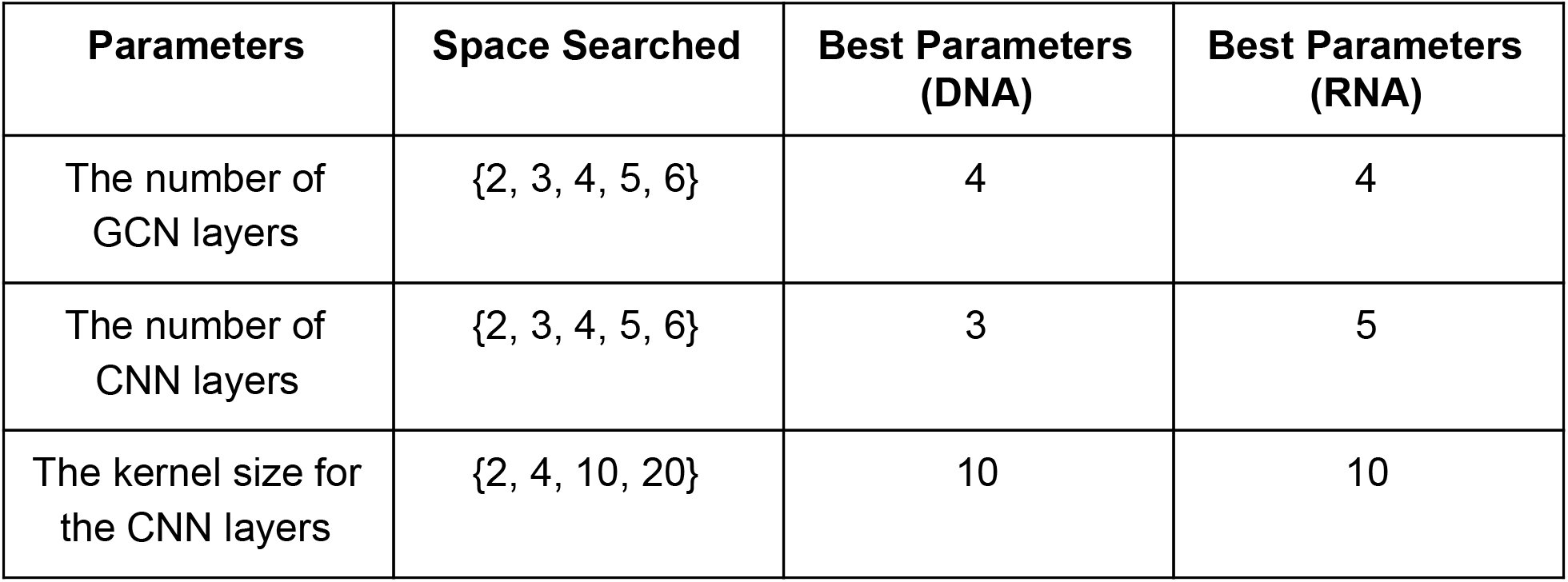
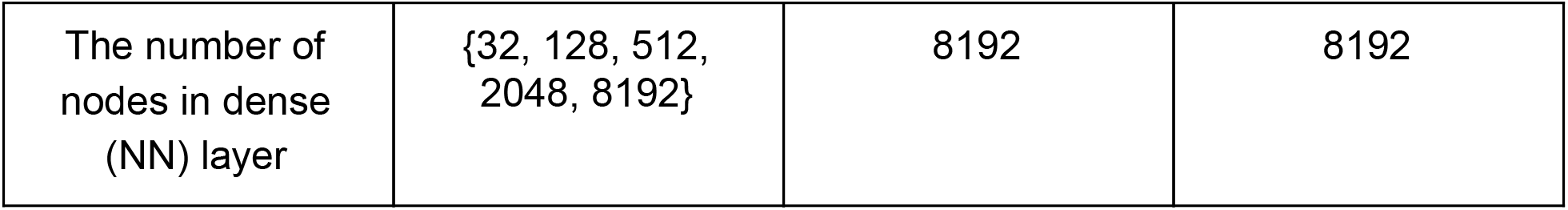

| Parameters | Space Searched | Best Parameters (DNA) | Best Parameters (RNA) |
| --- | --- | --- | --- |
| The number of GCN layers | {2, 3, 4, 5, 6} | 4 | 4 |
| The number of CNN layers | {2, 3, 4, 5, 6} | 3 | 5 |
| The kernel size for the CNN layers | {2, 4, 10, 20} | 10 | 10 |
| The number of nodes in dense (NN) layer | {32, 128, 512, 2048, 8192} | 8192 | 8192 |

We used the following scaling factor to determine the number of nodes in each GCN/CNN layer of our framework:

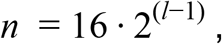

where *l* is the layer index of the GCN, CNN, and NN layer groups. For instance, the number of GCN layers determined to yield the best performance for DNA were 4. The number of nodes for each GCN layer was therefore 128, 64, 32, and 16. The same logic was applied to all other layer groups.

We performed 10-fold cross validation for each hyper-parameter combination. The combination that produced the lowest average RMSE across all folds was adopted as the optimal architecture. The optimal DNA framework has 4 GCN layers, 3 CNN layers with a kernel size of 10 and 8192 nodes in the NN layer. The optimal RNA framework has 4 GCN layers, 5 CNN layers with a kernel size of 10 and 8192 in the NN layer.

### Down-sample, base-dropout, position-dropout and combination analysis

For down-sample analysis, we performed random train-test splits in 5% intervals, noted as 0.95-0.05, etc. For base-dropout analysis, we created training sets by removing certain bases. Such train-test split creates 729/4096 (18%) training kmers and 3367/4096 (82%) test kmers for DNA, and 243/1024 (24%) training kmers and 781/1024 (76%) test kmers for RNA. It is important to note that everytime a base is dropped from the training set it is retained in the test set. Similar to base-dropout, the position-dropout adds one more dimension, which is the position of the nucleotide base. For a given position-dropout, the testing kmers are all kmers with the dropout nucleotide covering the target position, and the training kmers are the remaining kmers. Such position-dropout creates 3072/4096 (75%) training kmers and 1024/4096 (25%) test kmers for DNA, and 768/1024 (24%) training kmers and 256/1024 (25%) test kmers for RNA. It is important to note that bases dropped in a specific position in the training appear in the same position in testing. For combination analysis, we trained the framework by combining any of the two base-dropout kmer sets. For instance, all G and C-dropout DNA 6mers, which was noted as G-C. Such analysis creates 1394/4096 (34%) training kmers and 2702/4096 (66%) test kmers for DNA, and 454/1024 (44%) training kmers and 570/1024 (56%) test kmers for RNA. For each above-mentioned train-test split, in order to perform statistical analyses, we produced 50 independently trained frameworks for each experiment. Specifically, we performed 50-fold cross validation in the down-sample analysis, considering for each fold the train kmers were randomly selected. As for other analyses, we performed 50 independent repeats using the same training kmer sets. The variability among repeats came from the stochasticity of the training process. To confirm the robustness of our architecture, we further performed two independent replicates (Run-1 and Run-2) of 50.

### Predicting modification-containing DNA 6mers

For the 5mC imputation experiment, the framework was trained on all 4096 {A, T, C, G} DNA 6mers, plus {1%, 5%, 10%, 30%, 50%, 70%, 90%} of randomly selected 5mC-containing DNA 6mers, following the training process as described above. In order to perform statistical analyses, we produced 50 independently trained frameworks (50 independent repeats) for each category, with in total two independent replicates (Run-1 and Run-2) of 50. Such frameworks were then applied on all 15625 possible {A, T, C, G, 5mC} DNA 6mers.

For the chemical group-level generalization experiment, the framework was trained on all 4096 {A, T, C, G} DNA 6mers following the training process as described above. In order to perform statistical analyses, we produced 50 independently trained frameworks (50 independent repeats), with in total two independent replicates (Run-1 and Run-2) of 50. Such frameworks were then applied on all 15625 possible DNA 6mers, including those composed of {A, T, C, G, 5mC} and {A, T, C, G, 6mA}.

### Predictive analysis of predicted kmer models

#### Overview

To test whether the generated kmer models can be used to correctly interpret C-5mC status from nanopore readouts, we performed predictive analysis by using signalAlign to make per-read per-base predictions [2]. For a given reference position, signalAlign can produce posterior probabilities for all possible bases based on a provided kmer model. Thus, for DNA 6mer models generated as described in “predicting modification-containing DNA 6mers”, the empirical nanopolish [3,6] model obtained as described in “kmer models”, we allowed signalAlign to predict between C and 5mC. Considering no significant goodness-of-fit differences were observed between Run-1 and Run-2, only models generated in Run-1 were used here. All predictive analyses performed in this paper were within the human NA12878 cell line.

#### Selecting prediction sites

The prediction sites were selected among the entire human genome. To avoid artifacts caused by ambiguous genomic DNA modification status, we only focused on confident 5mC sites and canonical genomic regions in our analysis. Besides 5mC, other modifications exist in genomic DNA. Considering extremely low fractions of other modifications, e.g. only ~0.05% are modified as 6mAs in the human genome [23], we define “non-5mC” sites as “canonical regions” during predictive analysis. Among these canonical regions, we used the Poisson process with lambda equals 50 to randomly select genomic sites for signalAlign to predict. Such selected sites were at least 12 nucleotides apart, avoiding potential interference by the neighbors. We thus obtained confident 5mC and C sites for signalAlign prediction.

The genomic DNA C-5mC status was determined by analyzing two independent NA12878 cell line bisulfite sequencing datasets [24]. A C-site was determined as confidently methylated if, for both bisulfite sequencing datasets, 95% of reads were methylated with at least 10x coverage. On the other hand, a C-site was considered confidently unmodified if, for both bisulfite sequencing datasets, at most 1% of reads were methylated with at least 10x coverage. Such analysis covered 3367/3367 canonical C-containing DNA 6mers, and 3950/6144 single 5mC-containing DNA 6mers.

#### Selecting nanopore sequencing reads

We then ran signalAlign with reads reported in the nanopore consortium NA12878 cell line native genomic DNA datasets [25] covering the above-mentioned prediction sites. Considering the computational complexity of signalAlign, we performed the following filtering steps to use the fewest reads to cover the most kmers. First, we calculated read-level kmer coverage. For example, the center 5mC-site of DNA read CAGAT**<u>(5mC)</u>** ACAGA was selected for signalAlign prediction. 6mers CAGAT**<u>(5mC),</u>** AGAT**<u>(5mC)</u>** A, GAT**<u>(5mC)</u>** AC, AT**<u>(5mC)</u>** ACA, T**<u>(5mC)</u>** ACAG and **<u>(5mC)</u>** ACAGA span such 5mC-site, therefore considered as being covered. Based on such read-level kmer coverage, we iteratively selected reads that covered the least frequently covered kmers. Thus, building a read set which covers as many kmers as possible as often as possible with the fewest number of reads. We included two biological replicates of NA12878 cell line native genomic DNA sequencing experiments (FAB39088 and FAF01169) in the C-5mC predictive analysis. For such analysis, our final FAB39088 set contained 1706 reads, which covered 2625/3367 C-only DNA 6mers with an average 61.52x coverage as negative control, and 3105/3950 possible single-5mC DNA 6mers with an average 5.01x coverage. The final FAF01169 set contained 1396 reads, which covered 2610/3367 C-only DNA 6mers with an average 63.26x coverage as negative control, and 3140/3950 single-5mC DNA 6mers with an average 4.76x coverage. Combining the two sets, in total 2792/3367 C-only DNA 6mers were covered with an average 58.49x coverage, and 3481/3950 single-5mC DNA 6mers were covered with an average 4.38x coverage.

#### Performing signalAlign prediction

Based on the selected prediction sites and nanopore sequencing reads as described above, per-read per-site predictive analysis was performed by signalAlign. The signalAlign analysis was performed with default parameters, except for internal read-level quality filtering. Such quality filtering removes reads with poor kmer to ionic current correspondence. During signalAlign analysis, kmer-to-ionic current correspondence probability matrices (event tables) are first generated. Based on such event tables, signalAlign will remove reads with low average probabilities (<10^−5^). Additionally, reads with >50 consecutive ionic current signals that cannot be corresponded to kmers (probability equals 0) will be discarded. Considering the event table generation is based on the provided kmer model, therefore after the above-mentioned default quality filtering, the number of remaining reads varies when different kmer models are supplied during predictive analysis. To ensure the statistical soundness, we deactivate the default quality filtering, such that reads to be analyzed by different supplied kmer models will be the same.

#### Quantifying predictive accuracy

Based on the results of signalAlign prediction, we created confusion matrices (2×2 for 5mC predictive analysis with 5mC as “positive” class and C as “negative” class) to quantify predictive accuracy. Specifically, we calculated the true positive rate (TPR), true negative rate (TNR), positive predictive value (PPV), negative predictive value (NPV), F1-score (F1) and balanced accuracy (BA) as predictive accuracy quantifications. BA was presented in Figure 1E as representative quantification, and the full predictive performance can be found in Supplementary Figure 5.

## Kmer models

Canonical DNA 6mer and RNA 5mer models are available at: https://github.com/nanoporetech/kmer_models. The Nanopolish 5mC-containing DNA 6mer model is available at: https://github.com/nanoporetech/nanopolish/tree/master/etc/r9-models.

## GLOSSARY

Kmer: DNA or RNA sequence with length of k.
Canonical kmer: kmer sequences purely composed of non-modified nucleotides, including {A, T, G, C} for DNA and {A, U, G, C} for RNA.
Characteristic ionic current: ionic currents yielded by a specific kmer are usually modeled by a Gaussian distribution, the mean of which is referred to as the characteristic ionic current.
Kmer model: a table recording kmers and their corresponding nanopore sequencing characteristic ionic currents. To avoid confusion, the “deep learning model” will be referred to as “framework” throughout the paper.
Framework: in this paper “framework” specifically refers to the deep learning model used to predict the characteristic ionic current from kmer chemical structures.
GCN: Graph Convolutional Network.
CNN: Convolutional Neural Network.
NN: Neural Network.
RMSE: Root Mean Square Error.
R: Pearson correlation.
BA: Balanced accuracy.
5mC: 5-methylcytosine.
6mA: N6-methyladenine.
I: Inosine.
SMILES: Simplified Molecular Input Line Entry System for annotating chemical structures using character strings.
Atom: specifically refers to non-hydrogen atoms throughout the paper.

## Data availability

The FAB39088 and FAF01169 NA12878 cell line native genomic DNA nanopore sequencing datasets were downloaded from https://github.com/nanopore-wgs-consortium/NA12878/blob/master/Genome.md. The two independent NA12878 bisulfite datasets were downloaded from https://www.encodeproject.org/experiments/ENCSR890UQO/.

## Code availability

Codes for constructing, training and running the deep learning framework are available at https://github.com/ioannisa92/Nanopore_modification_inference. Codes for nanopore sequencing data analysis are available at https://github.com/adbailey4/functional_model_analysis. Codes for reproducing all figures are available upon request to the corresponding authors.

## ACKNOWLEDGEMENTS

Research reported in this publication was supported by the National Institutes of Health under Award Numbers R01-HG010053-02, U01HG010961, U41HG010972, R01HG010485, 2U41HG007234, 5U54HG007990, 5T32HG008345-04 and U01HL137183. The content is solely the responsibility of the authors and does not necessarily represent the official views of the National Institutes of Health. The authors would thank Jordan Eizenga, Dr. Jonas Sibbesen, Dr. Mark Akeson and Dr. Miten Jain for critical insight and help with drafting the manuscript.

## AUTHOR CONTRIBUTIONS

H.D. conceived the idea. I.A. performed deep learning framework modeling, optimization and analysis. A.B. and H.D. performed the nanopore sequencing data analysis. H.D., J.S. and B.P. supervised the project. All authors prepared the manuscript.

