## Supplementary for "Towards Inferring Nanopore Sequencing Ionic Currents from Nucleotide Chemical Structures"

### Supplementary Information

**Supplementary Figure 1. Goodness-of-fit of the canonical DNA analysis.** Root Mean Square Error (RMSE) and Pearson correlation (r) values of DNA down-sample, base-dropout, position-dropout and model combination analyses. Run-1 (solid boxes) and Run-2 (dashed boxes) refer to two independent replicates. RMSE and r values for the predictions of all DNA 6mers (Overall), DNA 6mers in training set only (Train) and DNA 6mers in test set only (Test) were marked as black, red and blue, respectively. See METHODS for details.

**Supplementary Figure 2. Goodness-of-fit of the canonical RNA analysis.** Root Mean Square Error (RMSE) and Pearson correlation (r) values of DNA down-sample, base-dropout, position-dropout and model combination analyses. Run-1 (solid boxes) and Run-2 (dashed boxes) refer to two independent replicates. RMSE and r values for the predictions of all DNA 6mers (Overall), DNA 6mers in training set only (Train) and DNA 6mers in test set only (Test) were marked as black, red and blue, respectively. See METHODS for details.

**Supplementary Figure 3. Goodness-of-fit of the DNA 5mC analysis.** Root Mean Square Error (RMSE) and Pearson correlation (r) values of DNA 5mC-imputation analysis. Run-1 (solid boxes) and Run-2 (dashed boxes) refer to two independent replicates. RMSE and r values for the predictions of all DNA 6mers (Overall), DNA 6mers in training set only (Train) and DNA 6mers in test set only (Test) were marked as black, red and blue, respectively. See METHODS for details.

**Supplementary Figure 4. RMSE correlation in DNA 5mC-*de novo* analysis.** For both Run-1 and Run-2, RMSE values obtained from canonical and 5mC-containing DNA 6mers were compared. Dots on the scatter-plots represent training-prediction repeats.

**Supplementary Figure 5. Predictive accuracy of DNA 5mC analysis.** Predictive accuracy was quantified by true positive rate (TPR), true negative rate (TNR), positive predictive value (PPV), negative predictive value (NPV), F1-score (F1) and balanced accuracy (BA). FAB39088 (black) and FAF01169 (red) refer to two independent NA12878 cell line native genomic DNA nanopore sequencing datasets [1]. See METHODS for details.

**Supplementary Figure 6. Visualizing canonical DNA 6mer atom similarity matrices.** Without losing generality, we visualized the atom similarity matrices of 10 random canonical DNA 6mers. Similarity matrices were calculated using the Pearson correlation of the state vectors outputted by the final GCN layers. Corresponding chemical structures of analyzed DNA 6mers were shown side-by-side of the similarity matrices, based on which atoms were numbered and colored. Carbon, nitrogen, oxygen and phosphorus were colored as black, blue, red and orange, respectively.

**Supplementary Figure 7. Visualizing 5mC-containing DNA 6mer atom similarity matrices.** Without losing generality, we visualized the atom similarity matrices of 10 random 5mC-containing DNA 6mers. 5mC was abbreviated as M for simplicity. Similarity matrices were calculated using the Pearson correlation of the state vectors outputted by the final GCN layers. Corresponding chemical structures of analyzed DNA 6mers were shown side-by-side of the similarity matrices, based on which atoms were numbered and colored. Carbon, nitrogen, oxygen and phosphorus were colored as black, blue, red and orange, respectively.

**Supplementary Figure 8. Chemical group stack analysis.** Framework trained with all possible canonical DNA 6mers was used to predict 6mA-containing 6mers. 6mA-containing kmers were grouped by the positions of 6mAs. Signal distributions of 6mA-containing kmers and their canonical counterparts were shown in the boxplot. See METHODS for details.

**Supplementary Note 1.** We first evaluated whether the proposed framework could generalize information to nucleotides that were not present in the entire training data. We thus trained the framework using the DNA 6mers that do not contain each nucleotide (base-dropout, see METHODS). Such training sets retain ~18% of the total 6mers. Therefore we used the 0.2-0.8 train-test split as the baseline null model. As shown in Figure 1B and Supplementary Figure 1, base-dropouts significantly decreased the prediction power compared to the baseline null model. Such a result suggests that the four DNA nucleotides provide orthogonal information during training. In addition, the prediction power was more impaired by excluding T and C, which suggests that the four nucleotides have unequal importance.

We also evaluated the framework's generalizability to nucleotides that were not present in particular DNA 6mer positions (position-dropout, see METHODS). Such position-dropout retains 75% of total 6mers for training, so we used the 0.75-0.25 train-test split as the baseline null model. As shown in Figure 1B and Supplementary Figure 1, in general the prediction power was significantly impaired by excluding T and C, consistent with the nucleotide importance evaluated by base-dropout analysis. Meanwhile, dropouts in 3rd and 4th positions contributed the most to prediction power decrease, followed by 2nd and 5th positions. The positional importance suggested here was further consistent with [2].

We further explored whether full DNA 6mer models can be generalized by combining complementary base-dropout training sets, e.g. G-dropout and C-dropout that contains instances of C and G containing kmers, but no kmers containing both C and G (noted as G-C, see METHODS). Such training sets contain ~34% of total DNA 6mers, thus 0.35-0.65 train-test split was used as the baseline null model. As shown in Figure 1B and Supplementary Figure 1, in general the prediction power was comparable with the baseline null model, suggesting the validity of such model combination.

**Supplementary Note 2.** Following the same pipeline as in DNA, down-sample, base-dropout, position-dropout and model combination analyses were also performed under RNA context. Meanwhile, RMSE and r were also used for prediction power evaluation for RNA analysis (see METHODS). Compared to DNA analysis, two major differences were observed. First, for RNA analysis as shown in Supplementary Figure 2, in general the prediction power was lower. For instance random down-sample analysis with 0.95-0.05 train-test split (best-performing random down-sample group), average RMSE values were ~0.8 and ~2.4 for DNA and RNA, respectively. We speculate that such prediction power difference was majorly caused by the number of training data points. As mentioned in the main text, with the currently most prevalent Oxford Nanopore Technologies R9.4 nanopore sequencing chemistry, DNA is modeled with in total 4096 6mers. On the other hand, RNA is modeled with in total 1024 5mers, only 25% as opposed to the DNA scenario. Such fewer possible training data points might strongly compromise the prediction power of our framework. However, once trained with a similar amount of kmers, the RNA architecture could yield comparable prediction power. For instance, the RNA 0.95-0.05 (972 training kmers) and DNA 0.25-0.75 (1024 training kmers) train-test splits yielded comparable performance. Such a result suggested the validity of our proposed architecture.

The other major difference between DNA and RNA analysis is, the four canonical DNA bases (A, T, G, C) are “orthogonal” to each other (Figure 1B and Supplementary Figure 1). In contrast, base-dropout will not cause statistically significant decrease in prediction power, suggesting that the four canonical RNA bases (A, U, G, C) can complement each other in terms of their chemical properties (Supplementary Figure 2). Here “orthogonal” means base-dropouts will significantly decrease the prediction power as opposed to the corresponding random down-sample null model. Notably, such an “orthogonality effect” was particularly strong for T and C. We speculate that such a difference can be explained by the additional methyl group in T. Among the four DNA canonical bases, methyl only appears in T, thus cannot be compensated by other combining A, G and C. Similarly, considering such methyl is encoded with pyrimidine backbone (Figure 2A and Supplementary Figure 6), the representation of the other pyrimidine nucleobase, C is also affected. Thus T and C were more “orthogonal” compared to A and G. As for RNA, without the additional methyl, the four canonical RNA bases complement each other in terms of their chemical structures. Further, the chemical information generalization among bases guarantees the proper representation of RNA 5mers under base-dropout scenario, thus producing statistical insignificant prediction power as opposed to the corresponding null model.

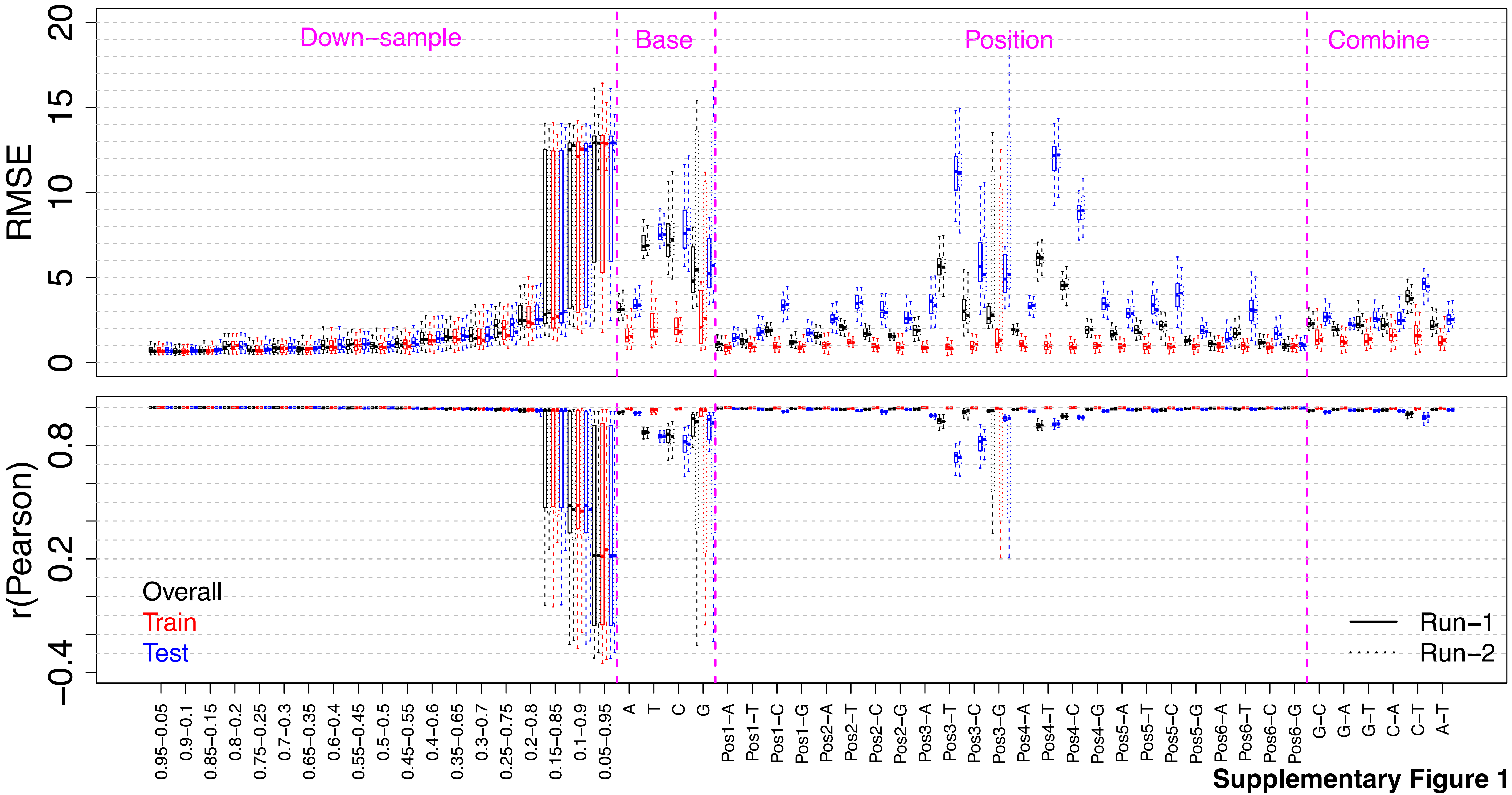

**Supplementary Figure 1**

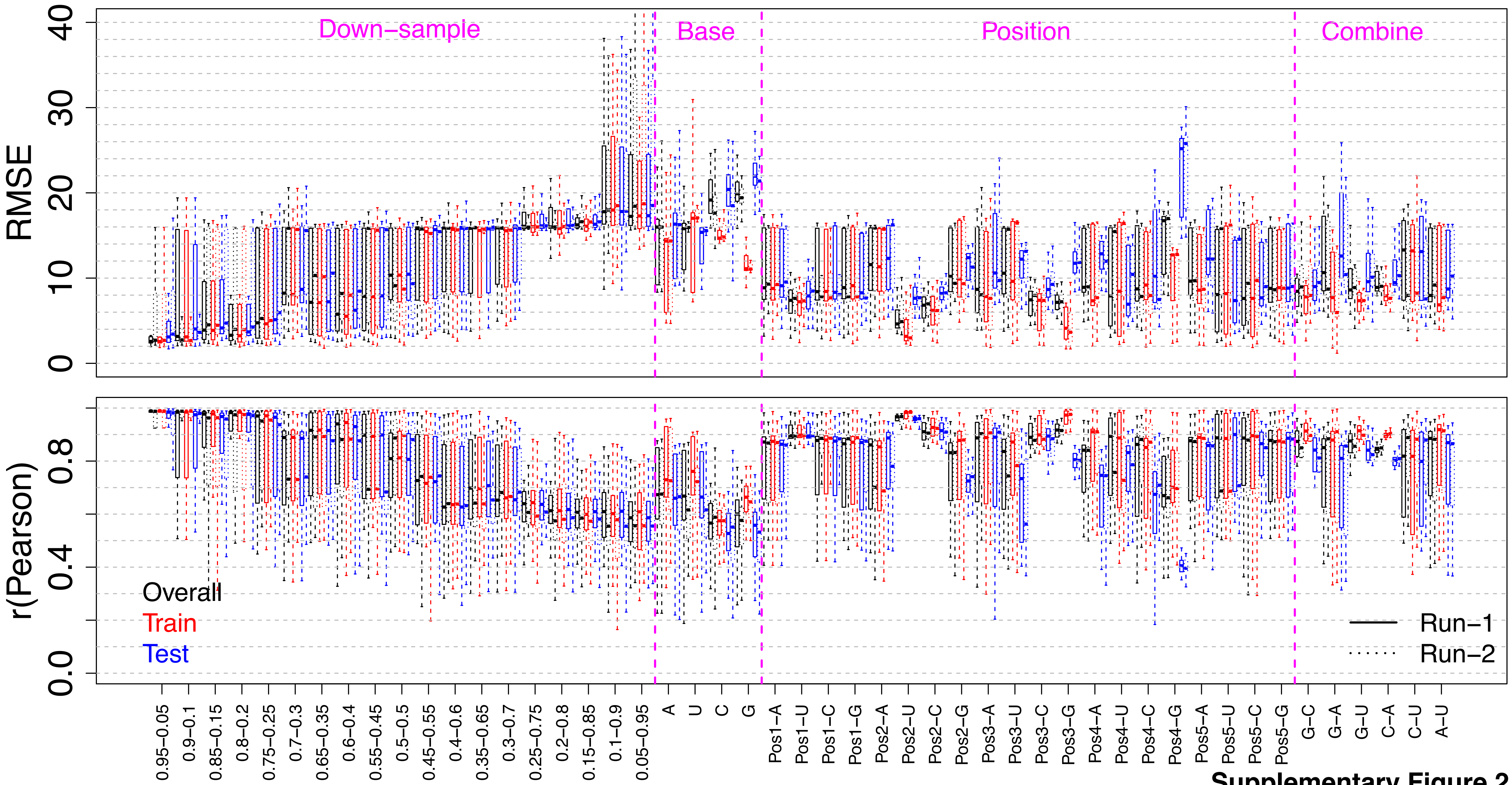


**Supplementary Figure 2**

**Supplementary Figure 3**

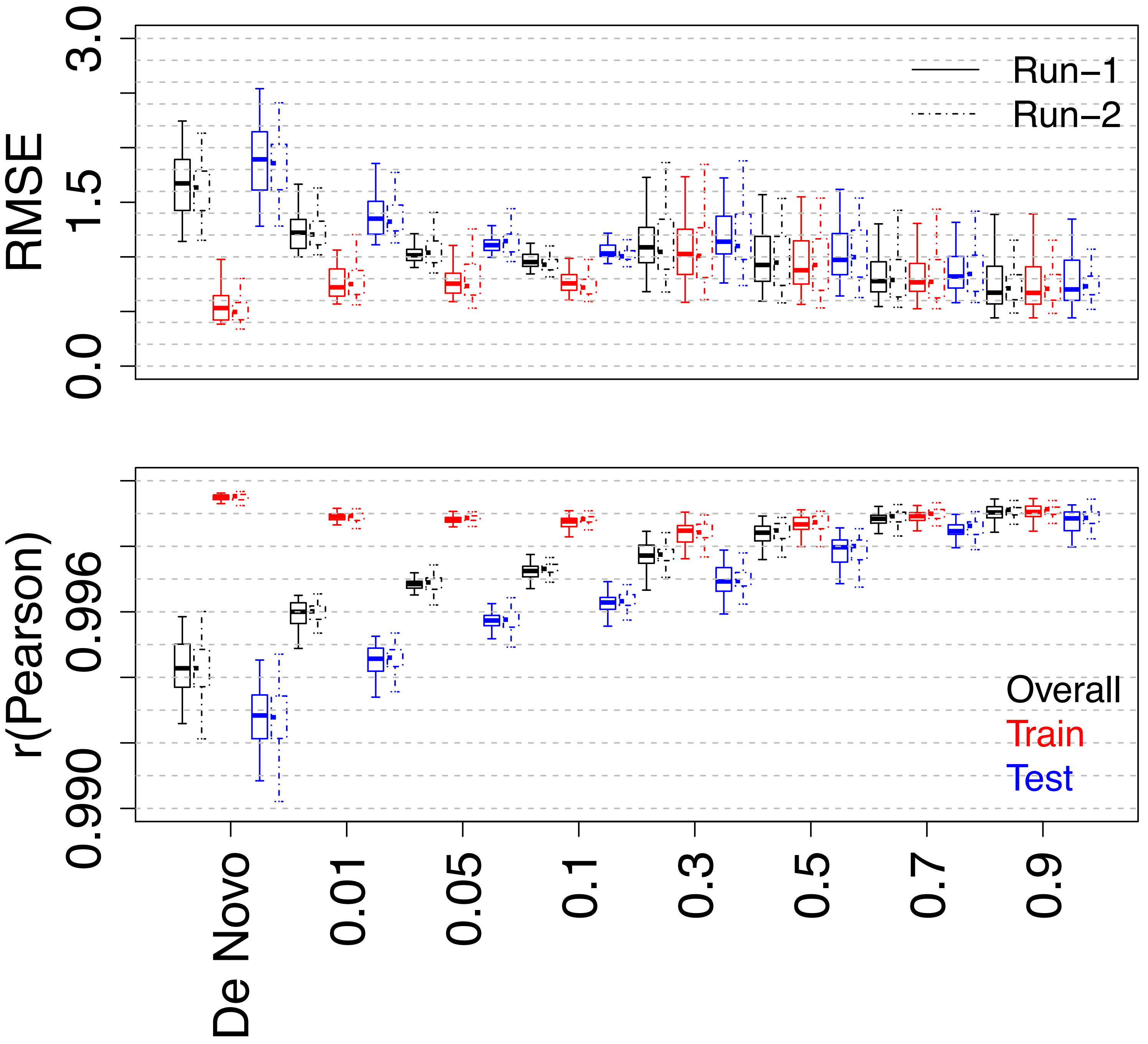

**Supplementary Figure 4**

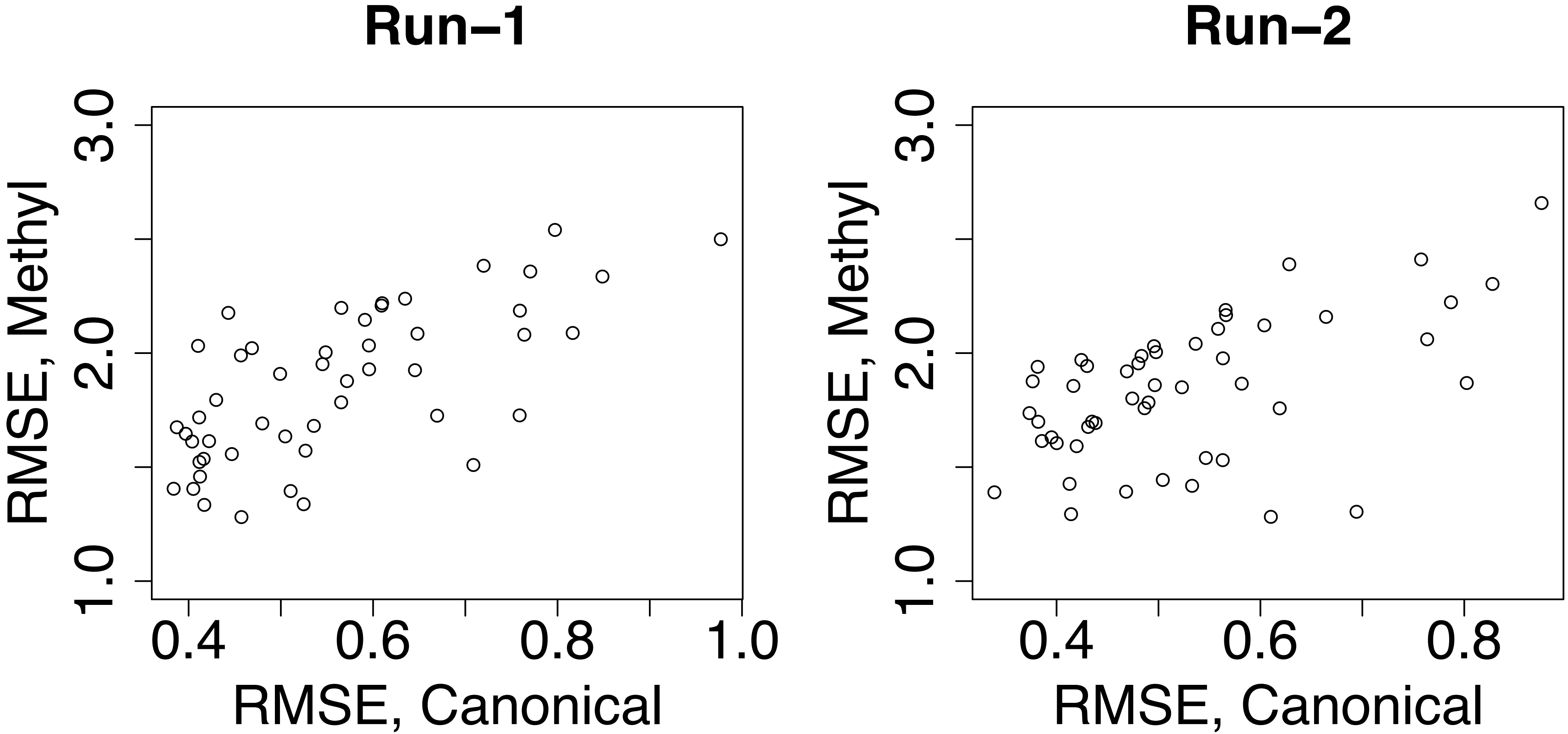

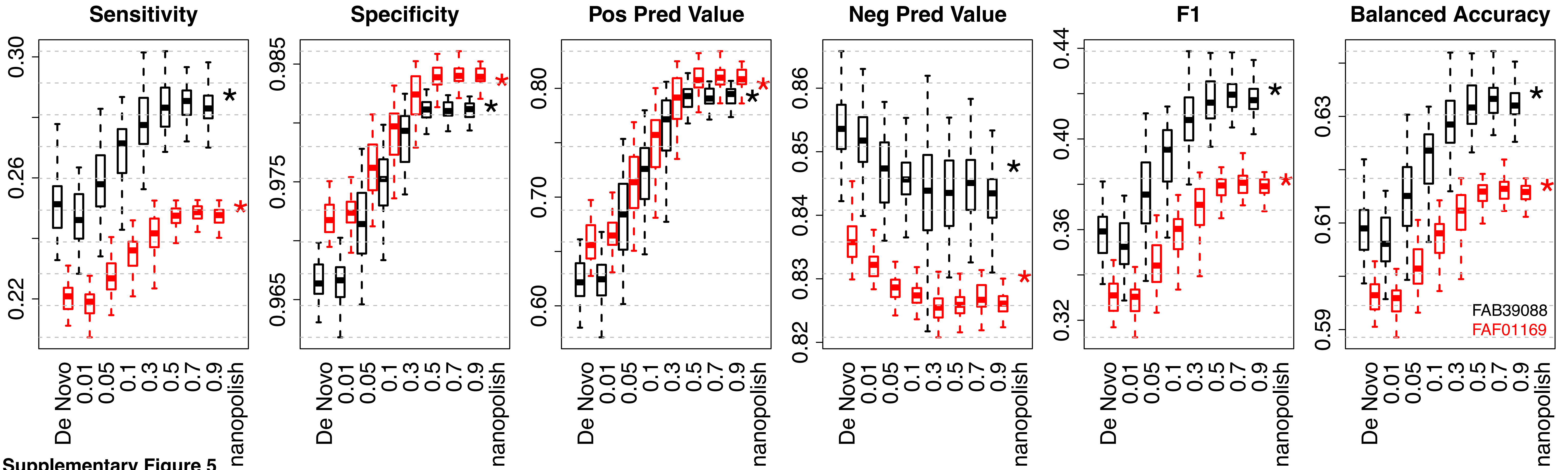


Supplementary Figure 5

Supplementary Figure 6

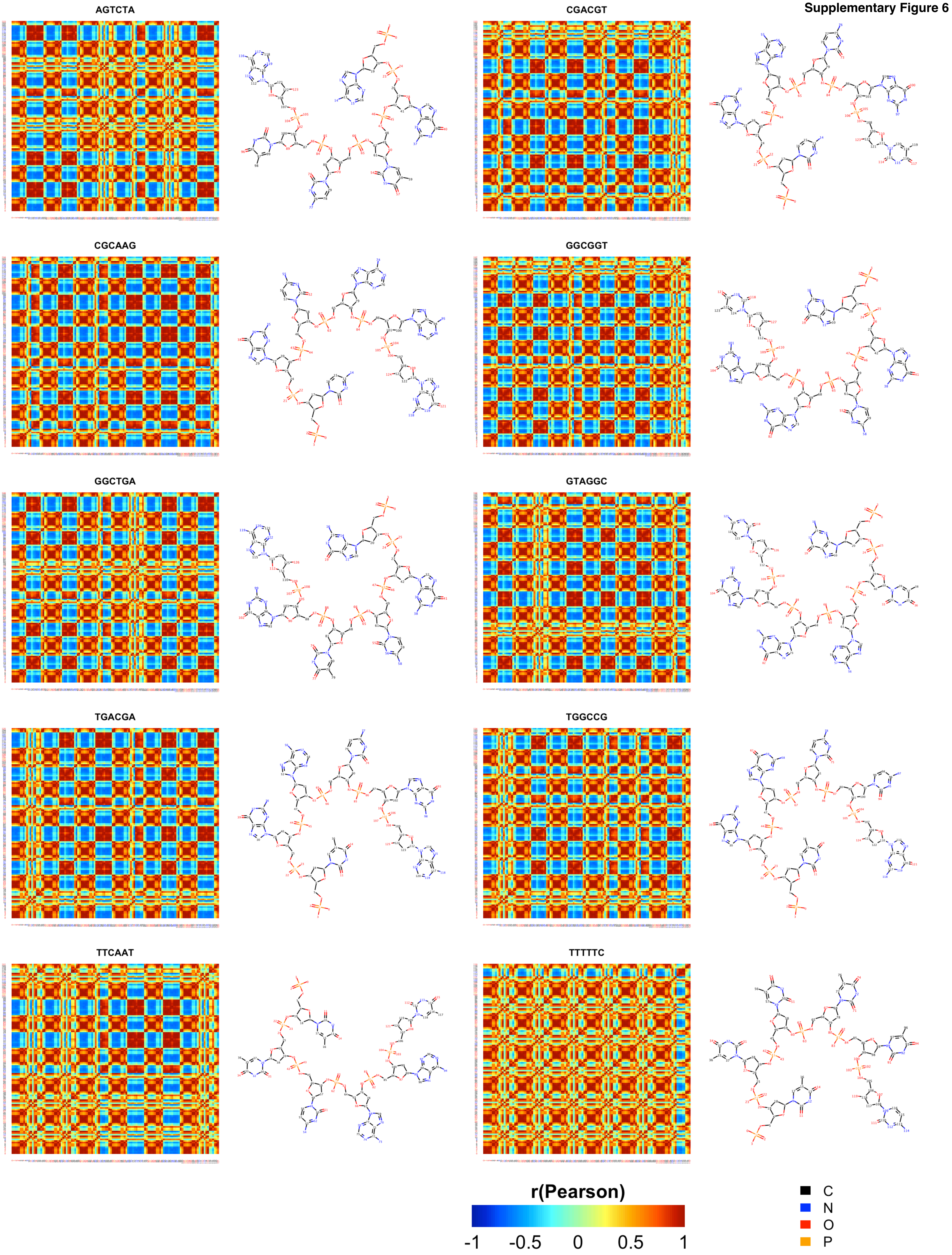

Supplementary Figure 7

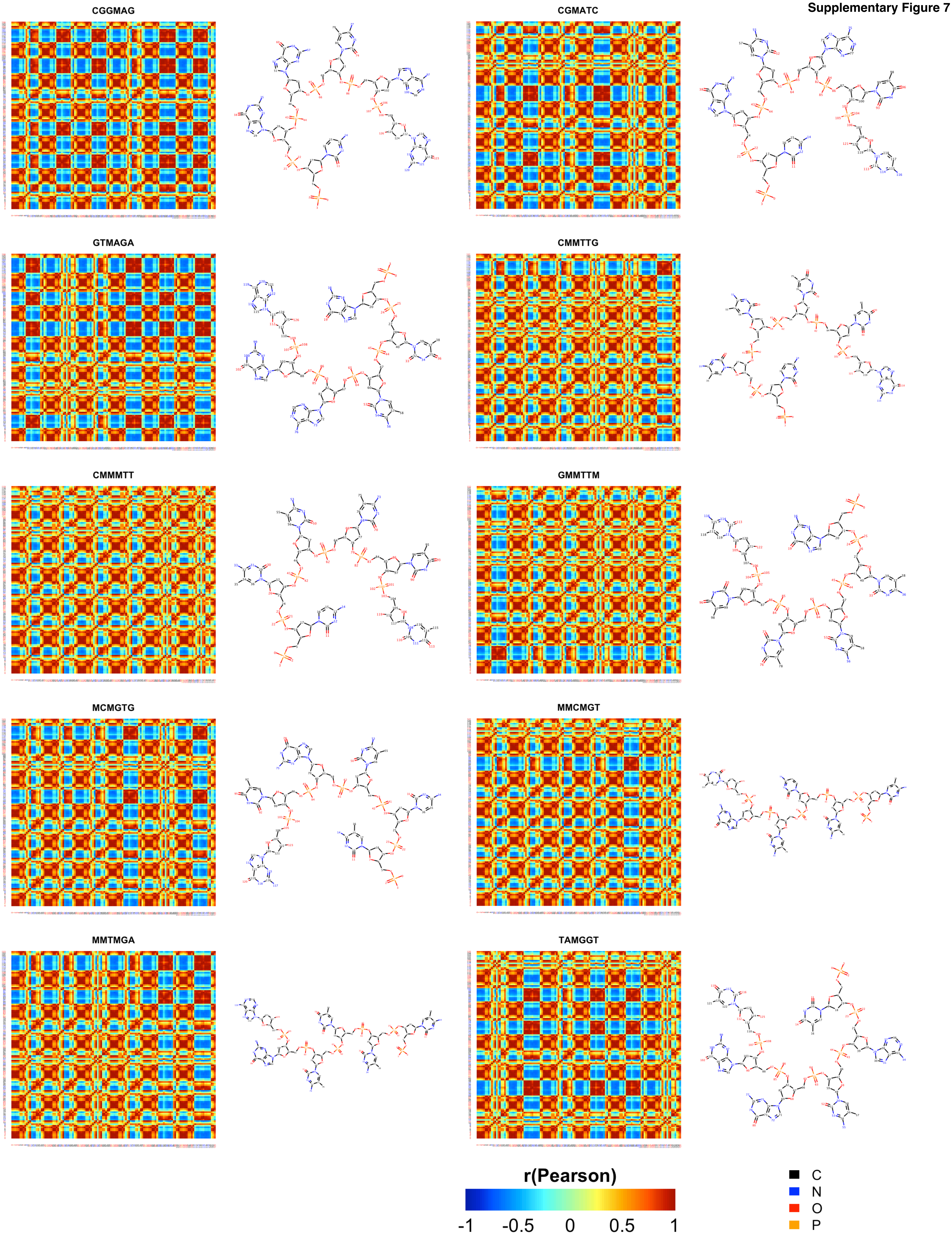

Supplementary Figure 8

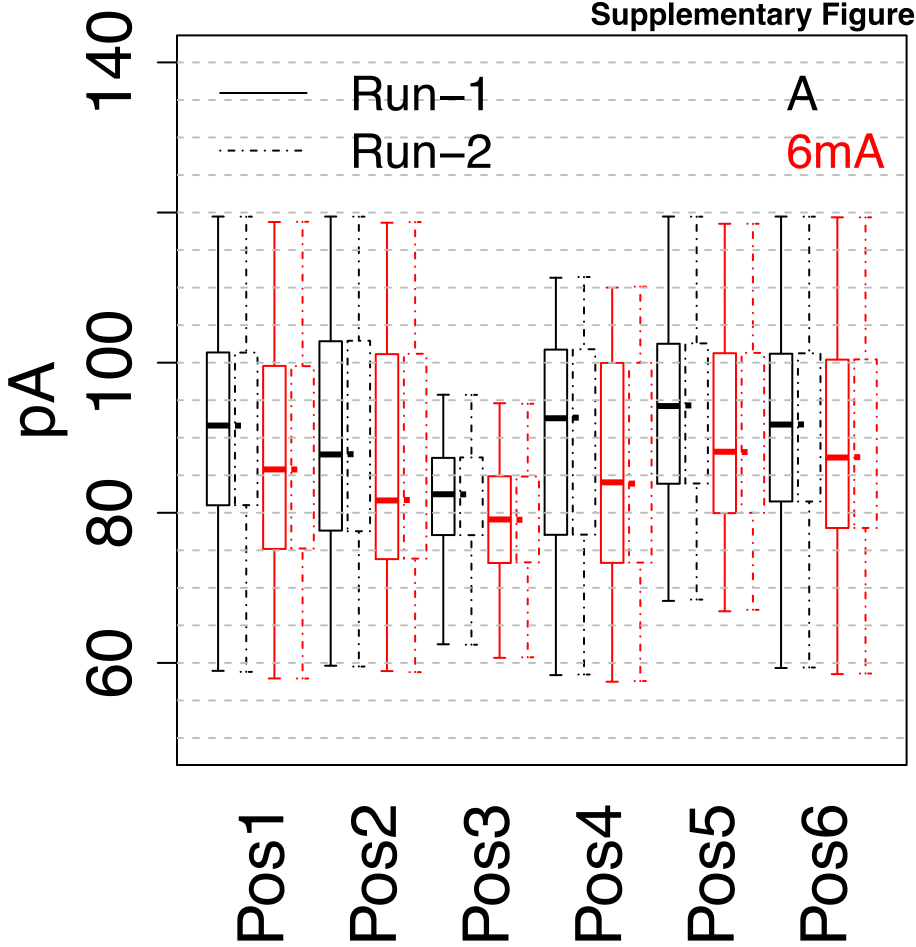
